# Measuring Regulatory Network Inheritance in Dividing Yeast Cells Using Ordinary Differential Equations

**DOI:** 10.1101/2024.11.23.624995

**Authors:** Wenbin Wu, Taylor Kennedy, Orlando Arguello-Miranda, Kevin Z. Lin

## Abstract

Quantifying the inheritance of protein regulation during asymmetric cell division remains a challenge due to the complexity of these systems and the lack of a formal mathematical definition. We introduce ODEinherit, a new statistical framework leveraging ordinary differential equations (ODEs) to measure how much a mother cell’s regulatory network is passed on to its daughters, addressing this gap. ODEinherit first estimates cell-specific regulatory networks through ODE systems, incorporating novel adjustments for non-oscillatory trajectories. Then, inheritance is quantified by evaluating how well a mother’s regulatory network explains its daughter’s trajectories. We demonstrate that precise quantification of this inheritance relies on pruning and adjustment for the network density. We benchmark ODEinherit on simulated data and apply it to live-cell, time-lapse microscopy data, where we track the expression dynamics of six proteins across 85 dividing *S. cerevisiae* cells over eight hours. Our results reveal substantial heterogeneity in inheritance rates among mother-daughter pairs, paving the way for applications in cellular stress response and cell-fate prediction studies across generations.

## 1 Introduction

Understanding how cells divide and pass on regulatory information across generations is fundamental to biology, with implications for processes such as cancer progression, immune responses, and tissue regeneration. Studying cell cycling, which is the process of how cells divide to yield new cells, unravels the potential for understanding tumor progression and cancer therapies (Otto and Sicinski, 2017; Ma and Gurkan-Cavusoglu, 2024). During cell cycling, the inheritance of molecular components can vary, especially in asymmetric divisions, where distinct daughter cell phenotypes emerge (Higuchi-Sanabria et al., 2014; Herrero et al., 2020). The *saccharomyces cerevisiae* (budding yeast), widely used to study cell cycling due to its rapid division rate, provides a model system in which mother cells can influence the fate of their daughters. Notably, studies such as those by Argüello-Miranda et al. (2018) have shown that a daughter’s fate can be predicted based on its mother’s properties, even before it is physically formed. These observations highlight the need to investigate how regulatory information shapes cell fate across generations.

Despite these advances, quantifying the inheritance of protein regulatory networks from mother to daughter cells remains challenging due to the dynamic and complex nature of these systems. Current approaches often fail to capture the network-level inheritance of cellular regulation. To address this gap, we propose a novel statistical framework based on live-cell, time-lapse microscopy data from yeast to rigorously measure the inheritance of protein regulatory networks. By bridging mathematical modeling with biological insights, our method quantifies the extent to which these networks are passed on from one generation to the next.

## 2 Background

### 2.1 Problem Setup

To investigate regulatory inheritance, we collected live-cell, time-lapse microscopy data tracking the cell division process, illustrated in Figure 1A. This data, which is the primary focus of this paper, simultaneously tracks the expression levels of six proteins in dividing yeast cells. Specific proteins were chosen a priori as markers of key cell cycle activities beforehand and were tagged with fluorescent reporters to visualize their quantities over time. We monitored 25 mother cells and their 60 daughter cells over eight hours, recording measurements every 12 minutes. This produced time series of protein expression (referred to as *trajectories* throughout the paper) for 60 mother-daughter cell pairs. While mother cells were observed continuously throughout the course, daughter cells were only observed after their birth. Further details on data collection and processing are provided in Appendix S1 and Ramakanth et al. (2024). Since yeast cells may begin in different cellular states, daughter cells may inherit different amounts of protein dynamics from their mothers. We first provide qualitative evidence for such heterogeneity. Figure 1B presents protein expression trajectories for two mother-daughter pairs, with each panel showing a different protein (out of the six measured proteins). In the red pair, the daughter cell visually follows the mother’s cyclical expression patterns after birth, suggesting a higher degree of inheritance. In contrast, the green pair displays notably weaker similarity in both the amplitude and frequency, indicating a lower level of inheritance. These examples highlight the possible heterogeneity in inheritance rates across the 60 cell pairs studied.

**Figure 1:**
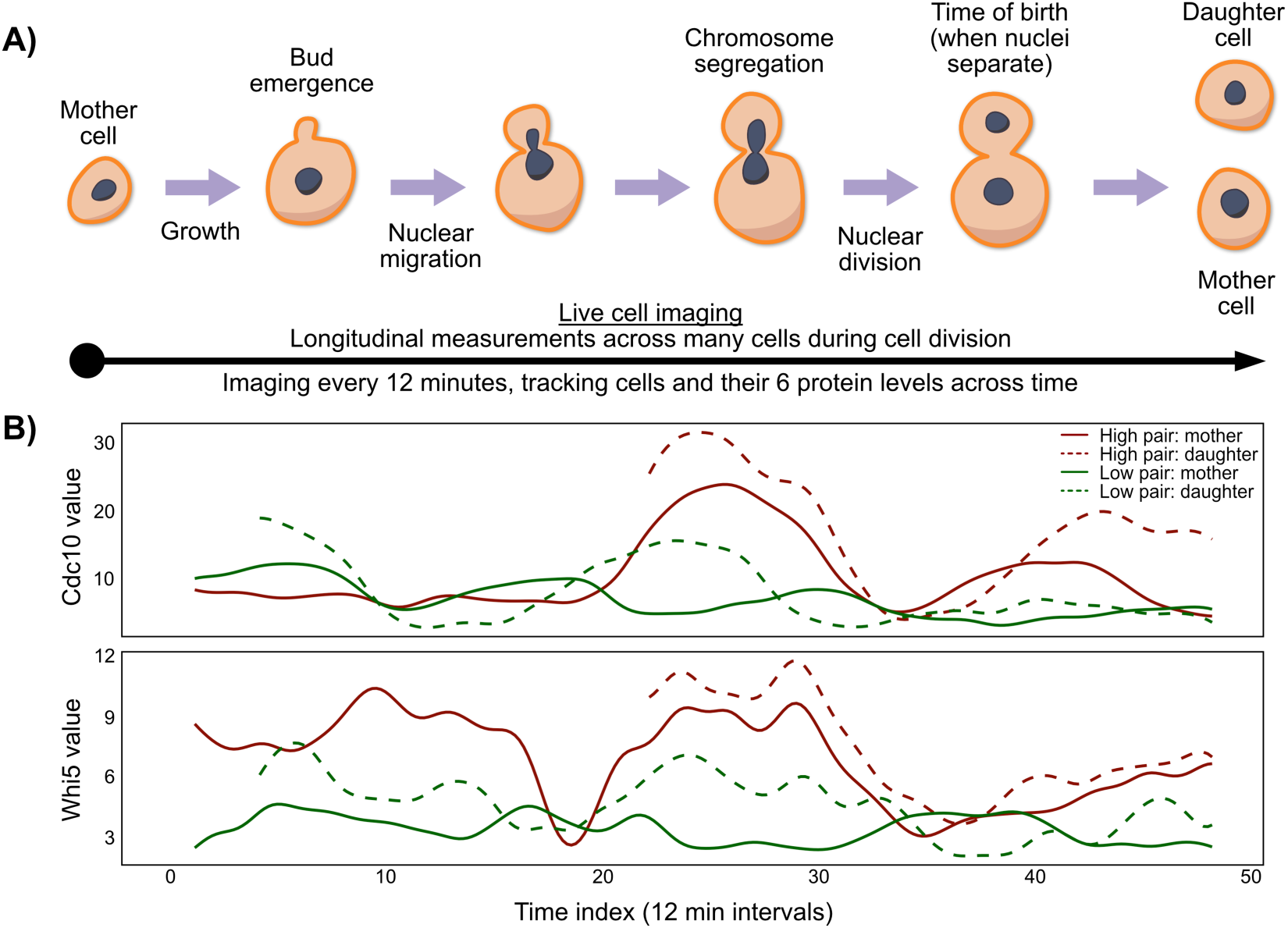
**A)**: Schematic representation of live-cell, time-lapse microscopy data collection during cell division. **B)**: Interpolated trajectories of two cell pairs with high (red) and low (green) degrees of inheritance, qualitatively speaking. For each pair, mother and daughter cells are shown with solid and dashed lines, respectively. Each panel shows the trajectory of one protein (out of six).

Our goal is to statistically quantify inheritance between mother and daughter cells. Although many statistical methods exist for analyzing time series data, their underlying premises may be biologically unrealistic in our context. For example, Granger causality tests whether the mother cell’s past protein expression can predict the daughter’s future expression. However, once separated, mother and daughter cells behave as distinct biological entities, and there is little biological reason to assume their protein levels should remain synchronized or predictive of each other. Instead, we hypothesize that inheritance occurs through the transmission of the mother cell’s underlying *regulatory machinery*, represented by the regulatory relationships between proteins (i.e., whether one protein regulates another). This hypothesis is motivated by the biological insight that, in asymmetric division, cellular components are unequally partitioned between mother and daughter cells. As a result, daughters inherit distinct initial protein abundances and regulatory states (Higuchi-Sanabria et al., 2014; Herrero et al., 2020). Inherited components, such as post-translational modifications and intermediary proteins, may influence how proteins interact within the daughter cell (Infant et al., 2021; Hamey and Wilkins, 2023). Thus, while the mother cell’s protein expression does not directly predict the daughter cell’s protein expression at any particular time point, the underlying regulatory structures may remain conserved between the two cells. Guided by this insight, we treat mother and daughter cells as separate entities and independently model their protein expression trajectories as functional data. Consequently, our statistical approach differs fundamentally from conventional time series analyses — we first model a mother cell’s regulatory network from its trajectories and then assess how much it explains the observed variability in the daughter trajectories.

### 2.2 Modeling as an Ordinary Differential Equation System

Mathematical modeling of protein regulations in cell cycles has long been a focus of research. Ordinary differential equation (ODE) systems have been widely used to model the temporal dynamics, particularly for their ability to capture feedback circuits that drive the oscillatory protein expression (Pomerening et al., 2003; Tyson and Novák, 2015; Sible and Tyson, 2007). In budding yeast, ODE models have been successfully applied to characterize protein regulatory networks (Chen et al., 2004; Radde and Kaderali, 2009; Boczko et al., 2010), including in our recent work on meiotic entry (Kociemba et al., 2024). Typically, each node in the network represents a protein, while directed edges indicate interactions such as activation, inhibition, or stabilization. However, estimating the parameters of an ODE system remains challenging – many of these aforementioned work studying yeast cells relies on manually selecting the parameters or performing a grid search (Tyson and Novák, 2015; Jashnsaz et al., 2021). In contrast, in this paper, we focus on estimating the ODE system directly from data.

Within the statistical literature, significant effort has been devoted to estimating ODE systems and studying properties of the resulting estimators (Dai and Li, 2022; Chen et al., 2017). Typically, the problem of modeling the interaction among p variables (i.e., proteins) is formulated as estimating a system of ODEs of the form,

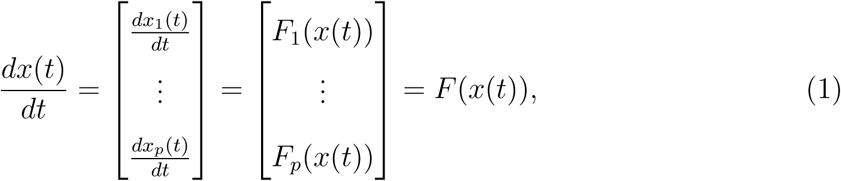

where the time index *t* is rescaled to the interval 𝒯 = [0, 1], and the unknown functionals *F* = {*F*_1_,..., *F_p_*} are the estimands that describe how variables regulate each other. In other words, this system models how the current states of all variables collectively influence the instantaneous rate of change of each variable. For a single sample (i.e., cell), the trajectories *x*(*t*) = (*x_p_*(*t*),..., *x_p_*(*t*)) ∈ ℛ*^p^* are observed at n discrete time points {*t*_1_, …, *t_n_*} with measurement errors,

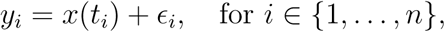

where *y_i_* = (*y*_*i*1_,..., *y_ip_*) ∈ ℛ*^p^* denotes the observed trajectories at time *t_i_*, and *ϵ_i_* = (*ϵ_i_*_1_,..., *ϵ_ip_*) ∈ ℛ*^p^* denotes the independent measurement errors with zero mean. We treat the initial conditions *x*(0) ∈ ℛ*^p^*as unknown.

Earlier studies primarily focused on estimating the regulatory functionals F*_j_* given prior knowledge of which variables regulate others. Various strategies have been developed to address the challenges of handling derivatives in the estimation process and ensure robust theoretical guarantees (Ramsay et al., 2007; Cao and Zhao, 2008; Liang and Wu, 2008; Brunel, 2008; Qi and Zhao, 2010; Xue et al., 2010; Gugushvili and Klaassen, 2012; Hall and Ma, 2014; Dattner and Klaassen, 2015). However, our study aims not only to estimate these functionals *F_j_* but also to identify the primary regulatory relationships among variables, which are naturally represented as a sparse regulatory network. This second goal adds complexity to the statistical task, requiring simultaneous estimation of the functionals and inference of the sparsity structure. Recent advances employing Lasso-type penalties have been introduced to encourage sparsity in the estimated functionals and to ensure selection consistency (Wu et al., 2014; Zhang et al., 2015; Chen et al., 2017; Dai and Li, 2022), making them especially relevant to our work. In this paper, we choose to build upon a particular nonparametric method called Kernel ODE (KODE, Dai and Li (2022)). This method is suitable as the backbone for our study since it leverages the flexibility of Reproducing Kernel Hilbert Space (RKHS) to capture the complex regulatory dynamics. Furthermore, the authors demonstrated its utility on *in silico* yeast data in their paper. We review further details in the next section.

## 3 Methods

In this section, we describe ODEinherit, our proposed method to measure how much a daughter cell inherits its mother’s regulatory machinery, which we will denote as *π*^(*M*^*^→D^*^)^. ODEinherit’s overall strategy involves first estimating a directed regulatory network for both mother and daughter cells by fitting an ODE system to their observed trajectories. Next, the inheritance score is defined as a percentage reflecting how well the mother network explains its daughter’s trajectories. We summarize this workflow in Figure 2. This procedure is applied to each mother-daughter pair to calculate the corresponding inheritance score. For simplicity, we omit the cell index in the notation and denote the observed trajectories of a given cell as 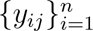 for *j* ∈ {1,..., *p*}.

**Figure 2:**
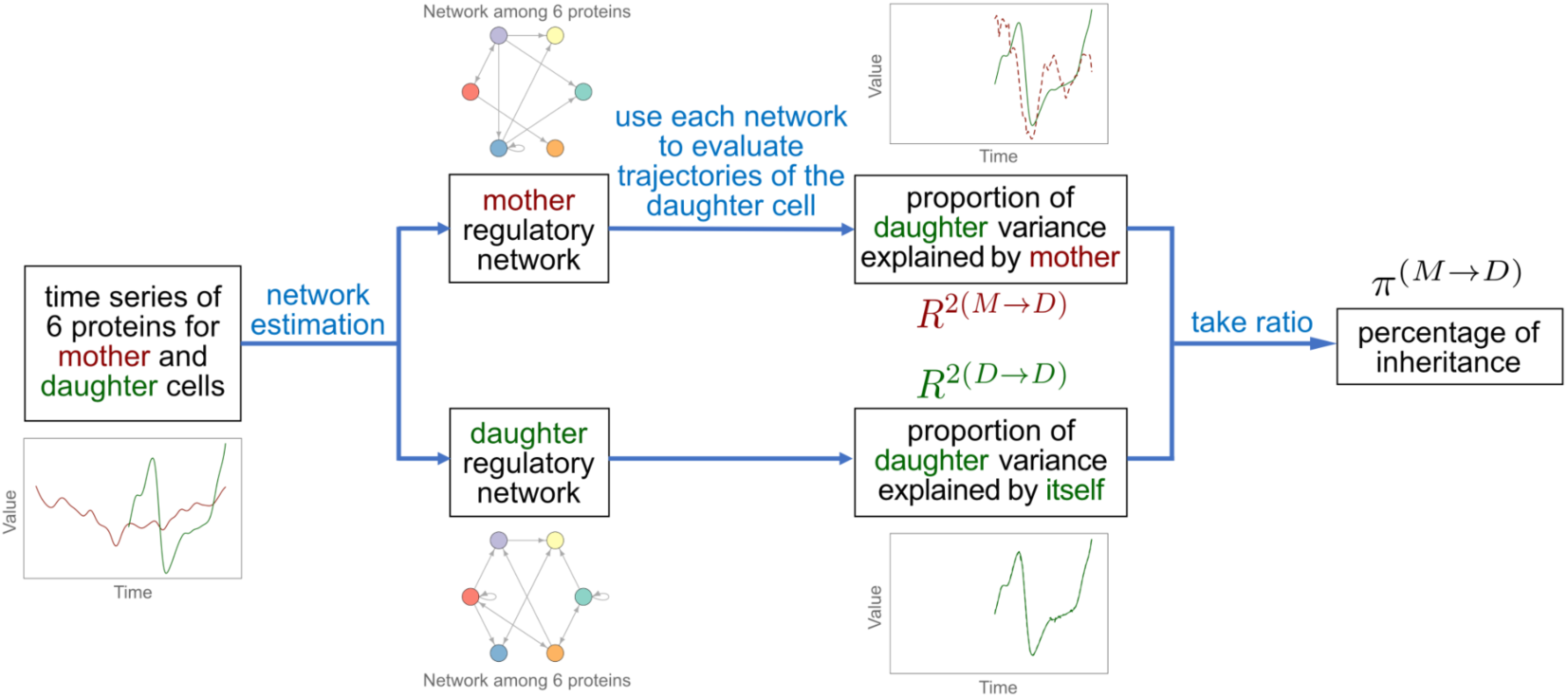
Workflow of measuring inheritance between a mother-daughter pair. In the example time series, the red and green time series refer to the mother and daughter cells, respectively.

### 3.1 ODE Estimation and a Review of Kernel ODE

We begin by describing how a directed regulatory network is constructed for each cell by estimating the ODE system in (1) from the observed trajectories. To model the regulatory relationships, we represent each functional *F_j_* using an additive structure, where the regulation of protein *j* is modeled as the sum of effects from all variables:

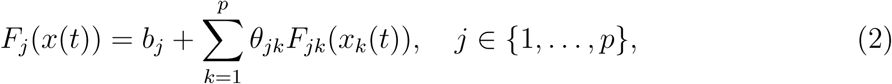

where *b_j_* ∈ ℛ is the intercept, *θ_jk_* ∈ ℛ and *F_jk_* characterizes the coefficient and dependency of variable *j* on each variable *k*, respectively. If *θ_jk_* ≠ 0, we consider variable k to be a regulator of variable *j* and assign a directed edge from variable *k* to *j* to construct the regulatory network.

We review the estimation framework of Dai and Li (2022) (KODE) here, as our method will build upon this framework for our inheritance analysis. For each variable *j*, KODE assumes that the derivative functional *F_j_* resides in the RKHS 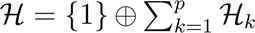, where 𝓗*_k_* is the RKHS generated by a given Mercer kernel *K_k_* corresponding to variable *k* such that *F_jk_* ∈ 𝓗*_k_*. In KODE, *F_j_*is estimated using a two-step collocation strategy, which is known to be computationally effective for ODE estimation. The first step obtains smoothing estimates to the observed trajectories by solving

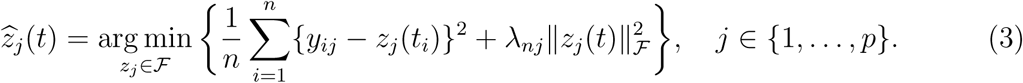

where 𝓕 is a given space of smooth functions. We denote the smoothing estimates as 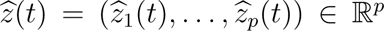. The second step estimates each functional *F_j_* ∈ 𝓗, the coefficients θ*j* = (θ_*j*1_,..., θ*_jk_*) ∈ ℛ^*p*^, and the initial condition θ_*j*0_ = *X_j_*(0) ∈ ℛ by solving the following penalized optimization problem,

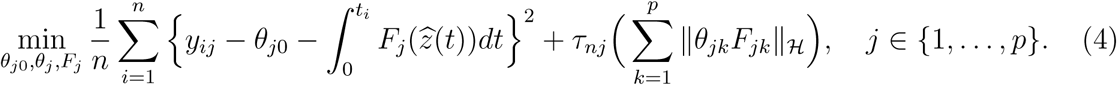

An iterative optimization algorithm is used for estimation, where sparsity in the estimated coefficients 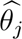 is induced by a Lasso regularization. Letting 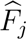 and 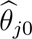 be the solution from (4), we recover the trajectories by

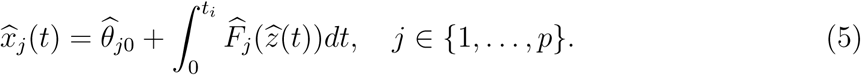

The integration is evaluated using a first-order approximation over a fine grid on 𝓣. The estimated regulatory network is constructed using the estimates 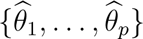. We use 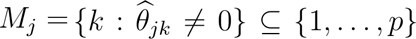 to denote the selected regulators of each variable *j*, then the estimated directed network is represented by {*M*_1_,..., *M_p_*}. We review additional details of KODE, such as data-driven strategies to tune hyperparameters, in Appendix S2.

### 3.2 Measuring the Goodness-of-Fit of a Network

We now introduce a statistic that measures the goodness-of-fit of the directed network represented by {*M*_1_,..., *M_p_*} based on how well it recovers the observed trajectories. This statistic plays a central role in our definition of inheritance in subsequent sections. For each variable *j*, we apply the KODE estimation algorithm to refit 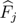 based on the smoothing estimates 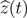 by removing the sparsity regularization and restricting *θ_j_* ∈ *M_j_*. Let 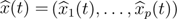 be the recovered trajectories from (5). We obtain the following variable- specific *R*^2^ statistic,

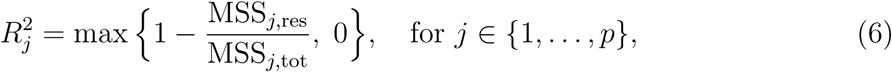

where 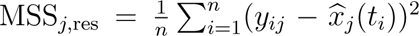 and 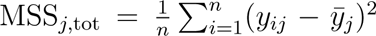 are the mean sums of squares of the residuals and of the observations, respectively, and 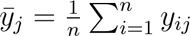 Intuitively, *R_j_*^2^ measures the proportion of variance in variable *j*’s observations that is explained by the recovered trajectory 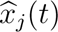. As shown in Equation (6), in cases where a negative *R_j_*^2^ value arises, the mean of the observations provides a better fit than our estimation, and we set *R_j_*^2^ = 0. To obtain an overall goodness-of-fit statistic for the network, we use

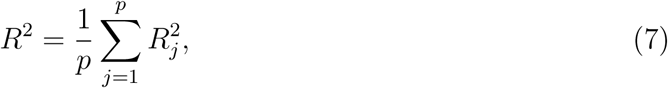

namely, the average of *R_j_*^2^ over all variables.

Unlike previous studies that primarily focus on the asymptotics of the estimation errors such as 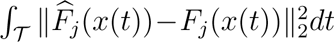 (Dai and Li, 2022), we use our *R*^2^ statistic as a goodness-of-fit measure for the estimated network at the individual-cell level. This is because, in these previous studies, there was no necessity to compute *R*^2^ at the individual-cell level. Notably, refitting a cell’s trajectories using its own network results in *R*^2^ values near 1 if the true *F_j_* resides in the correct model space. In contrast, we define our goodness-of-fit metric (7) to serve two key purposes: (1) quantifying inheritance by assessing how well the mother’s regulatory network explains the daughter’s trajectories, and (2) sparsify the network by evaluating the explanatory power of each selected regulator as we will describe in Section 3.4. As we will discuss in the following sections, both roles are central to our analysis of regulatory inheritance.

### 3.3 Measuring Inheritance Between Mother and Daughter Cells

We are now ready to describe how we quantify the daughter cell’s inheritance of the mother’s regulatory machinery. Our proposed score uses the *R*^2^ statistic to evaluate how much the mother’s regulatory network explains the daughter trajectories. Consider a mother-daughter pair whose estimated regulatory networks are denoted by *M* (mother) and *D* (daughter). We fit the daughter trajectories twice — once using the mother network M, and once using the daughter’s own network *D* — yielding two *R*^2^ values, *R*^2(*M→D*)^ and *R*^2(*D→D*)^. We take the ratio of these two values to define a percentage of inheritance for the mother-daughter pair:

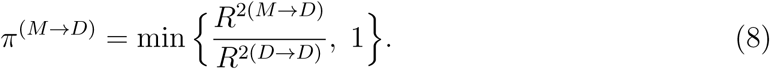

Here, *R*^2(*D→D*)^ measures how well the daughter’s own network explains its trajectories, reflecting intrinsic explainability, while *R*^2(*M→D*)^ measures how well the mother network explains these same trajectories. The quantity *π*^(*M→D*)^ ∈ [0, 1], which we call the *inheritance score*, is the primary output of ODEinherit, with 0 indicating no inheritance and 1 indicating full inheritance of the mother’s regulatory machinery.

The biological rationale behind this score is that the regulatory functionals *F_j_*’s are not transmitted identically from mother to daughter cells. Instead, they undergo modifications as the daughter cell progresses independently through its cell cycle, limiting the ability of the mother’s regulatory functionals to directly predict the daughter trajectories. Our inheritance score circumvents this limitation by measuring how much of the daughter’s variability can still be explained when its model space is restricted to the mother’s regulator sets. Thus, our score captures the inheritance of regulatory machinery without requiring the functionals themselves to remain unchanged across generations.

### 3.4 Pruning the Network

In this section, we describe a key issue we empirically observed that can adversely impact the inheritance score and describe our procedure for mitigating it. Consider the definition of our inheritance score *π*^(*M→D*)^ in (8). Similar to the regression setting, the explained variability by a model naturally increases as the number of covariates (i.e., regulators in the mother network) grows. Hence, as the mother network *M* becomes denser, the numerator *R*^2(*M→D*)^ tends to approach 1. This leads to inflated inheritance scores regardless of the daughter’s trajectories, which is an undesirable quality since the score becomes uninformative about the mother-daughter relations.

Our empirical findings indicate that KODE is more prone to producing false positives (i.e., selecting variables as regulators when they are not) than missing true regulators (i.e., false negatives), despite already using a Lasso-like penalty. This leads to overly dense estimated networks. As we demonstrate in later simulations, such excessive density severely impairs our ability to measure regulatory inheritance. To address this issue, we introduce a pruning strategy that refines and sparsifies each network while only slightly compromising its goodness-of-fit. Our strategy treats the regulator sets selected by KODE as initial candidates and iteratively prunes them based on their contribution to the trajectory recovery, quantified using the *R*^2^ statistic. Specifically, for each variable *j*, we evaluate the impact of removing each selected regulator by measuring the change in *R*^2^. A regulator is pruned if its removal reduces *R_j_*^2^ by less than a pre-specified threshold (by default, 5%). The detailed steps of this pruning procedure are provided in Appendix S2. This approach significantly reduces the false positive rate at a cost of slightly increasing the false negative rate. Overall, this results in sparser and more interpretable networks while ensuring no important regulators are omitted. We provide a further discussion on this pruning procedure in Appendix S2.

## 4 Simulation Study

In this section, we describe a suite of simulations to assess the performance of our network estimation, even in misspecified settings, and demonstrate that ODEinherit can meaningfully estimate the regulatory inheritance between simulated mother and daughter cells.

### 4.1 Network Estimation

We first evaluate the empirical performance of the network estimation strategy for both additive and non-additive ODE systems, where the assumed additive form (2) is violated in the latter case.

#### 4.1.1 Additive ODE System

We first demonstrate the accuracy of our network estimation procedure on a simple linear ODE system where the true regulatory functionals *F_j_*’s follow the assumed additive structure (2). This simulated system consists of two triplets of variables with solution trajectories being combinations of sine and cosine functions. It is specified as follows: for triplet *k* ∈ {1, 2}

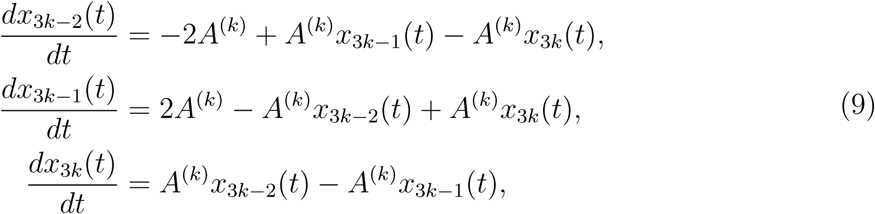

for time *t* ∈ [0, 1]. Here, *A*^(1)^ and *A*^(2)^ are chosen to make the triplets exhibit 3 and 10 periods on the interval [0, 1], respectively, minimizing the correlation between the two triplets. We simulate *n* = 200 time points at the evenly-spaced time grid {1/*n*, 2/*n*, . . ., 1}, using identical and independent Gaussian errors 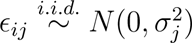.

Figure 3A demonstrates that our network estimation is accurate after pruning. Specifically, we plot the ROC curves for the estimated networks before and after pruning, where the noise variance *ς_j_*^2^ is set to 0% (i.e., no noise), 3%, and 10% of the sample variance of the true trajectory values 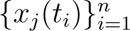 for each variable *j* ∈ {1,..., 6}. For each noise level, we perform 100 simulation runs and calculate the false positive rate (FPR) and true positive rate (TPR) for each run. The ROC curve and the area under the curve (AUC) are computed using the FPR and TPR. The results show that, in the presence of noise, the estimated networks without pruning tend to be overly dense, yielding high FPRs. In contrast, pruning substantially reduces the FPR, resulting in sparser and more interpretable networks. In the Appendix S3, we also show that the pruned networks maintain a similar level of explanatory power for the observed trajectories, with *R*^2^ values remaining close to those obtained before pruning.

**Figure 3:**
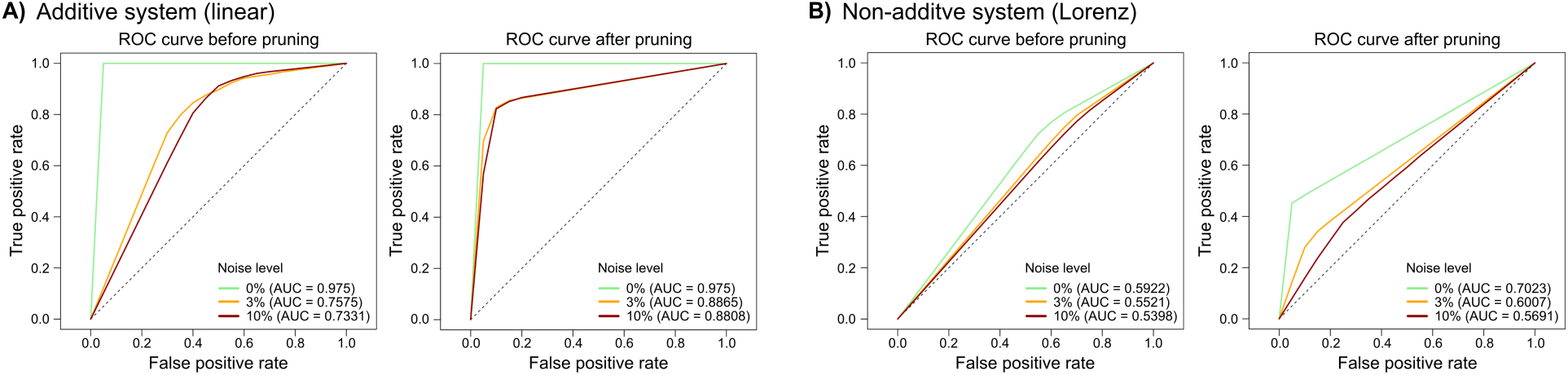
ROC curves of the network estimation in the additive **(A)** and non-additive system **(B)** under various noise levels, before (left) and after (right) our pruning procedure. The noise level is defined as the ratio of the noise variance, *ω*^2^, to the variance of the true trajectory *x_j_*(*t*). The black dashed line represents the performance of a random classifier (baseline *AUC* = 0.5).

#### 4.1.2 Non-Additive ODE System

We next demonstrate that we can still estimate meaningful networks from more complex trajectories when the true generative model is not within the assumed model space. In this simulation, we use a non-additive ODE system where the additivity assumption (2) is explicitly violated. Specifically, we consider the Lorenz system, a non-linear and aperiodic ODE system with interaction terms. The generating ODE model includes two triplets from this system with different sets of parameters: for triplet *k* ∈ {1, 2},

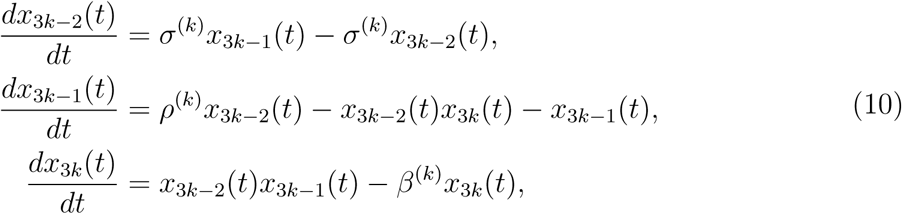

for time *t* ∈ [0, 100]. We set the parameters 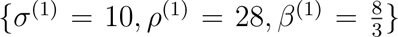 and 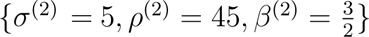 such that the trajectories oscillate chaotically. We simulate *n* = 200 time points at an evenly-spaced time grid over the intervals [40, 50] and [40, 60] for the first and the second triplets, respectively, and then rescale the time intervals to [0, 1]. This ensures that the frequencies are different for the triplets and are low enough for the oscillations to be captured in the n = 200 time points. The measurement errors are 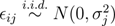, where *ς_j_*’s reflect the noise levels as described in Section 4.1.1.

Despite analyzing misspecified data, our results demonstrate that pruning enhances the accuracy of our estimated networks. For simplicity, an edge is assigned in the regulatory network whenever one variable affects another either through an additive or interaction effect. For example, we assign an edge *x*_3*k*_ → *x*_3*k−1*_ if *x*_3*k*_ affects *x_3*k−1*_ via an interaction with *x*_3*k*−2_*. Figure 3B displays the ROC curves for the estimated networks. Although model misspecification reduces overall selection accuracy, the pruning procedure still substantially improves performance while maintaining a similar level of explanatory power for the observed trajectories. See Appendix S3 for *R*^2^ comparisons before and after pruning.

### 4.2 Inheritance Score

With a reliable network estimate, we now move on to evaluate ODEinherit’s ability to reliably measure the amount of protein regulatory inheritance. To do this, we need to extend our simulation framework in order to generate mother-daughter pairs under each of the two ODE systems in Section 4.1. Our new simulation framework involves two components: a mother system and a pseudo-daughter system, which are used to generate the trajectories of the observed mother cell and an unobserved “pseudo-daughter” cell, respectively. The actual daughter trajectories are then generated as a convex combination of these two trajectories where the combination weight controls the degree of inheritance between the mother and daughter cells. Intuitively, the pseudo-daughter trajectories represent the daughter’s intrinsic dynamics without inheritance, while the actual daughter trajectories reflect partial inheritance from the mother.

We provide an overview of how we simulate mother-daughter pairs. Consider an ODE model with two triplets, either (9) or (10). In the mother system, we set the triplets *j* ∈ {1, 2, 3} and *j* ∈ {4, 5, 6} to adhere to the generative model as before. However, in the pseudo-daughter system, variables 2 and 5 are interchanged, resulting in the triplets *j* ∈ {1, 5, 3} and *j* ∈ {4, 2, 6} following the same dynamics. The corresponding networks are shown in Figure 4. The mother cell trajectories, denoted by *x*^(*M*)^(*t*) ∈ ℛ^6^, are generated from the mother system over n = 200 evenly-spaced time points on the standardized interval [0, 1] at a noise level of 1%. The daughter cell is assumed to be born at time *t* = 0.3. The pseudo-daughter trajectories, denoted as *x*^(*D,*pseudo)^(*t*) ∈ ℛ^6^, are generated on the interval [0.3, 1] with initial conditions *x*^(*D,*pseudo)^(0.3) set to *x*^(*M*)^(0.3) at its time of birth. Let *α* denote the weight of mother trajectories, with larger values indicating higher degrees of inheritance. The actual daughter trajectories *x*^(*D*)^(*t*) ∈ ℛ^6^ are then simulated as

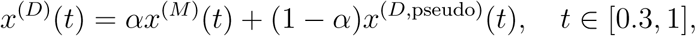

**Figure 4:**
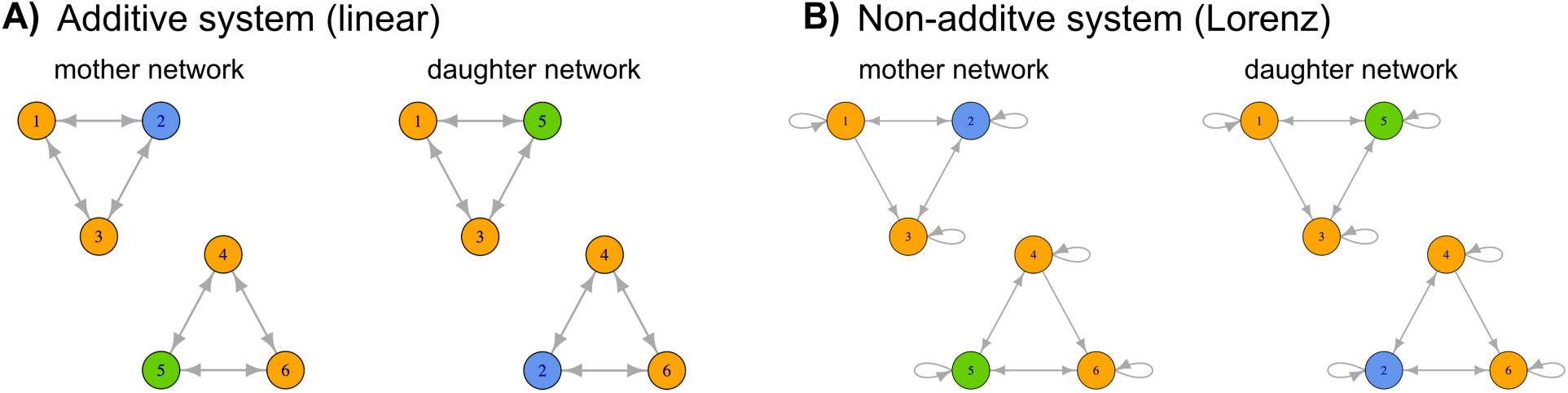
True mother and daughter regulatory networks in the additive system (**A**) and the non- additive system (**B**). Variables 2 and 5 are interchanged for the daughter, such that its network has an equivalent density as the mother network.

where data are drawn on the same observation time grid, restricted to the interval [0.3, 1], and subject to measurement errors at the 1% level.

To assess whether ODEinherit captures the amount of inheritance accurately, we create a set of simulated mother-daughter pairs with varying degrees of inheritance. Specifically, we generate *R* = 20 mother-daughter pairs for each *α* ∈ {0, 0.1, 0.2,..., 1}, as described above for both the additive and non-additive systems. Here, *α* = 0 and *α* = 1 denote no or full inheritance, respectively, and we are interested to see if ODEinherit can measure meaningful differences in inheritance between these two extremes. We employ the estimation workflow in Section 3.3 and obtain the inheritance score *π*^(*M→D*)^ for each cell. We note here that *α* does not directly represent the percentage of inheritance, and therefore *π*^(*M→D*)^ is not a direct estimate of *α*. Additionally, due to the nature of our ODE estimation approach, even a random network can account for a nonzero portion of the variance in the observed trajectories. In an extreme case when no regulation exists at all (i.e., an empty network), the fitted functionals 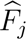’s in (2) become constants, and the recovered trajectories are simply linear trends. These trends might still explain some variability in the daughter trajectories, yielding a positive *π*^(*M→D*)^ inheritance score even when no inheritance exists. To account for this, we establish a baseline for how much a random mother network can explain the daughter trajectories. We do this by generating 10 random mother networks of equivalent density for each estimated mother network. These random networks are then used to refit the daughter trajectories and calculate the corresponding inheritance scores. Figure 5 shows that ODEinherit’s regulatory inheritance score is strongly correlated with *α*, the degree of inheritance dictated in our simulation. This plot shows the inheritance score *π*^(*M→D*)^ against *α* for each system, using the original, pruned, and random networks. We make three remarks about the results. First, the *π*^(*M→D*)^ estimate generally increases as *α* grows, indicating higher inheritance when the mother cell has more influence on the daughter cell. Notably, when there is no inheritance (i.e., *α* = 0), the *π*^(*M→D*)^ scores are indistinguishable from those using random mother networks, validating ODEinherit’s ability to detect a lack of inheritance. Second, even under model misspecification, the *π*^(*M→D*)^ estimate maintains an increasing trend with *α* but exhibits greater variability due to the reduced accuracy in network estimation. In this misspecified setting, the baseline yields higher *π*^(*M→D*)^ values because the Lorenz system has a denser true network, which inherently explains more variability in the trajectories. Third, without pruning, the *π*^(*M→D*)^ scores are largely inflated due to the high FPR in network estimation. The false positive edges inflate the R^2(*M*^*^→D^*^)^ values, which leads to inflated *π*^(*M→D*)^ scores. This issue becomes particularly severe in complex ODE systems under model misspecification, where the *π*^(*M→D*)^ scores are all close to 1 regardless of the dictated inheritance level *α*, thereby obscuring meaningful inheritance quantification. The pruning procedure in ODEinherit is essential for correcting this inflation and improving the accuracy of the inheritance score ε^(*M*∈*D*)^.

**Figure 5:**
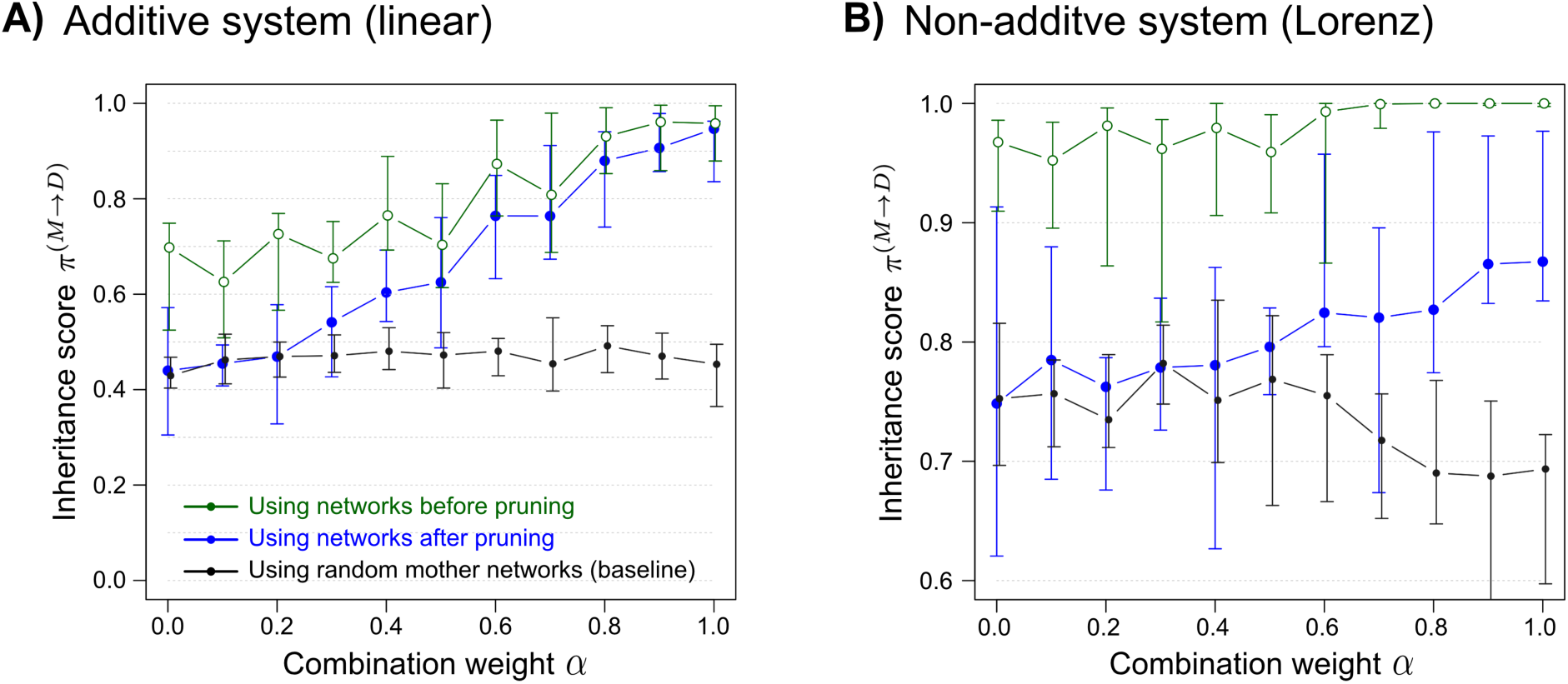
The inheritance score *π*^(*M→D*)^ against the weight of mother trajectories (*α*) in the additive (**A**) and non-additive (**B**) systems, respectively. Larger *θ* values indicate higher degrees of inheritance. Points indicate the median scores across the simulated pairs, and error bars represent the first and third quartiles.

## 5 Investigation of Inheritance in Yeast

We now return to the motivating data analysis described in Section 1 and demonstrate how ODEinherit enhances our understanding of cellular dynamics.

### 5.1 Data Details and Preprocessing

We first describe the data and preprocessing steps before presenting the results of ODEinherit. The dataset consists of 85 cells observed over 48 time points, including 25 mother cells and 60 first-generation daughter cells. Our analysis focuses exclusively on mother cells and these first-generation daughters, as these daughters have longer trajectories compared to second- or third-generation daughters. The longer trajectories offer more samples for accurate estimation. To filter out noise, we used a combination of Functional Principal Component Analysis (FPCA) and local polynomial regression to simultaneously smooth and interpolate the trajectories, achieving a fivefold temporal resolution relative to the original time grid. Additional details are provided in Appendix S1.

### 5.2 Adjusting Inheritance Score by Mother Network Density

We applied ODEinherit to each mother-daughter pair to infer their regulatory networks and calculate the corresponding inheritance score. Since the true generating model is unknown, we used the first-order Matéern kernel with the same implementation as in Section 4.1.2, as it provides a more flexible function space to capture the complexity of the cell trajectories. Before passing into KODE, we removed the linear trends in each cell’s trajectories.

While our pruning procedure sparsifies the estimated networks, cells still naturally exhibit varying numbers of regulatory edges (i.e., network density). As discussed in Section 3.4, differences in mother network density can influence the degree to which mother networks explain daughter trajectories, complicating the direct comparisons of inheritance scores across cell pairs. Indeed, the left panel of Figure 6A shows a positive correlation between the inheritance score and mother network density (Pearson correlation *ρ* = 0.24, p-value = 0.07). To adjust for this, we performed a resampling procedure: for each mother-daughter pair, we generated 400 randomized mother networks by randomly selecting regulators for each variable while preserving the original network density. We then normalized the original inheritance score by subtracting the mean and dividing by the standard deviation of the resampled scores, yielding the *adjusted inheritance score*. This adjustment parallels the baseline comparisons using randomized networks used in our simulation studies. As shown in the right panel of Figure 6A, the adjusted inheritance score no longer correlates with mother network density (*ρ* = 0.03, p-value = 0.8), enabling more comparable measurements of regulatory inheritance on the population level.

**Figure 6:**
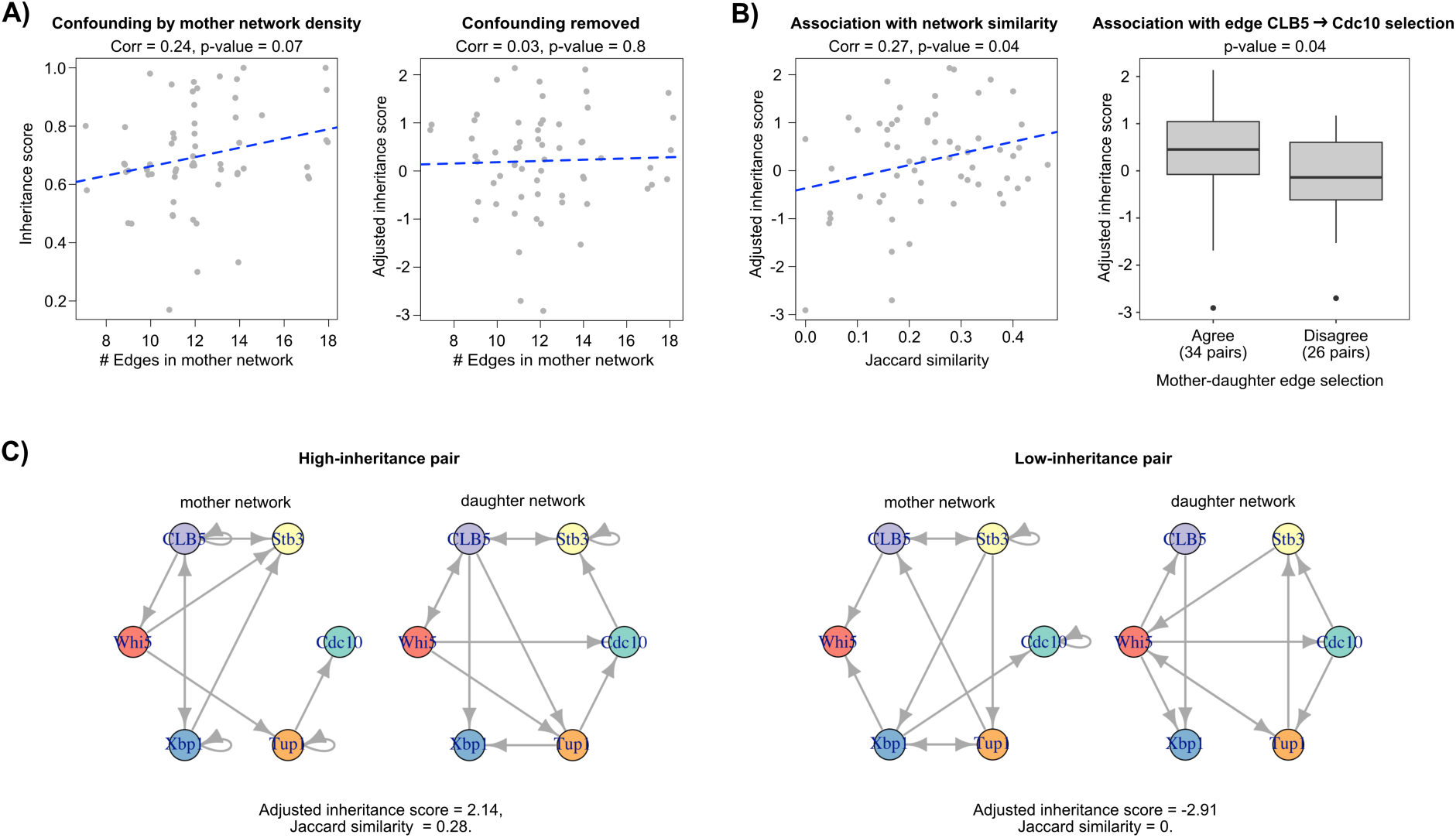
Inheritance score in yeast cells. **A)**: Confounding of inheritance scores by mother network density (left), corrected after normalization (right). Each point represents a mother- daughter pair. **B)**: Relationship between adjusted inheritance scores and network similarity (left), and adjusted inheritance scores stratified by agreement on the edge CLB5 → Cdc10 (right). Higher Jaccard similarities indicate more similar mother-daughter network structures. “Agree” denotes pairs where both mother and daughter either selected or did not select the edge (34 pairs), and “Disagree” denotes pairs where their selection differed (26 pairs). **C)**: Example networks illustrating high (left pair) and low (right pair) inheritance scores in mother-daughter pairs.

### 5.3 Association With Network Similarity

We now investigate whether the inheritance score is associated with similarity in the network structure. We quantify similarity between the mother and daughter networks, M and D, using the Jaccard similarity, defined as

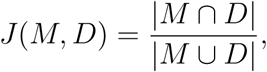

where |*M* ⋂ *D*| is the number of regulatory edges shared by both networks, and |*M* ⋂ *D*| is the total number of unique edges present in either network. As shown in Figure 6B (left), we find a positive correlation between the Jaccard similarity and the adjusted inheritance score (*ρ* = 0.27, p-value = 0.04). This relationship supports the validity of our inheritance score, as a similar network structure is typically associated with stronger inheritance. To further validate our approach, we focus on the regulatory edge from CLB5 to Cdc10, which is supported by biological evidence (Segal et al., 1998). In Figure 6B (right), mother-daughter pairs are stratified by whether they agreed (both selected or both did not select) or disagreed (only one selected) on the edge CLB5 → Cdc10. Pairs that agreed on this edge exhibited significantly higher inheritance scores (p-value = 0.04 using t-test), providing biologically grounded support for our approach. We also demonstrate two examples of mother-daughter network pairs in Figure 6C, one with high inheritance and one with low inheritance.

### 5.4 Cell Trajectories Alone Do Not Predict Regulatory Inheritance

Lastly, we investigate whether mother cell trajectories can provide insight into their regulatory networks and inheritance to their daughters. Based on the primary cell cycle activity marker Cdc10, we again applied FPCA to all mother cells and grouped them into two main clusters using K-means clustering on the first five FPCA components. Figure 7A depicts the trajectories of Cdc10 for each cluster. It is seen that cells in Cluster 1 exhibit fewer oscillations with larger amplitude, whereas cells in Cluster 2 undergo multiple cell cycles with greater consistency. This raises the question: do trajectory patterns of the mother cells alone inform the amount of regulatory relationships the daughter cell will inherit?

**Figure 7:**
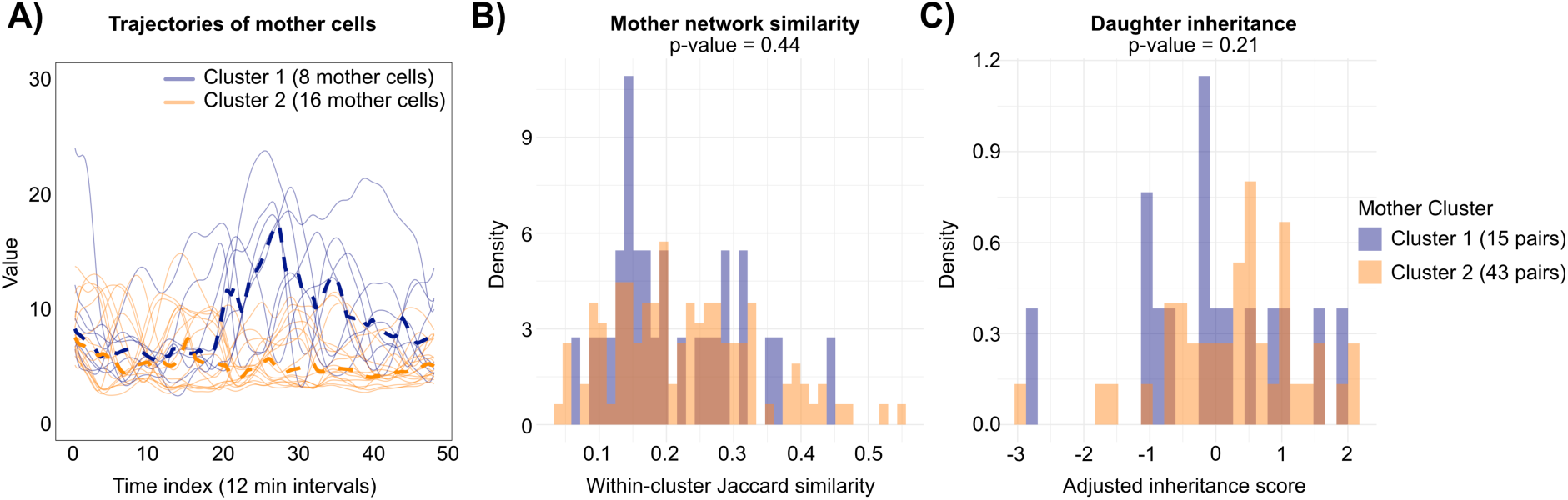
A) Interpolated trajectories of the primary cell cycle marker (Cdc10) for mother cells in each cluster; dashed lines indicate cluster medians. **B)** Pairwise Jaccard similarities between mother networks within each cluster; p-value from Wilcoxon rank-sum test comparing clusters. **C)** Adjusted inheritance scores of mother-daughter pairs by cluster; p-value from Wilcoxon rank-sum test comparing clusters.

Figure 7B presents the distribution of pairwise Jaccard similarities between mother networks within each cluster. A Wilcoxon rank-sum test comparing the clusters yields a p-value of 0.44, indicating no significant difference. Figure 7C shows the distribution of adjusted inheritance scores for mother-daughter pairs in each cluster. Although Cluster 2 (more cyclic cells) shows a slight increase in inheritance scores compared to Cluster 1, the difference is not statistically significant (p-value = 0.21). These results suggest that cell trajectories alone carry limited information about regulatory network structures and inheritance. In contrast, ODEinherit extracts these higher-order properties, revealing heterogeneity across cells that cannot be inferred by examining protein trajectories alone.

## 6 Conclusion

Understanding how regulatory information is transmitted across generations during cell division is a fundamental question in cell biology. Building on the hypothesis that a daughter cell inherits its regulatory machinery from its mother, we developed a novel statistical framework to quantify the extent of this inheritance. Our approach involves estimating an ODE system to model protein regulatory dynamics and measuring inheritance based on how effectively the mother regulatory network predicts the daughter trajectories. To ensure reliable results, our inheritance score depends on a sparse regulatory network, which is achieved through a heuristic pruning procedure. Simulations validated the effectiveness of our method. We employed the method to investigate budding yeast cells and revealed heterogeneity in inheritance rates across lineages, highlighting its potential to uncover biologically meaningful insights.

In this work, we primarily focused on ancestor cells and their first-generation daughter cells, as their longer observed trajectories provide more reliable data for modeling. Future research could explore how heritability influences cell fates across multiple generations, potentially uncovering long-term developmental trends in cell populations, as suggested by studies like Mura et al. (2019). However, as later generations often contain less observations, additional statistical considerations would be needed to address these challenges. Another promising direction is to extend our framework to study cellular responses to experimental stimuli, leveraging ODE methods that have been adapted for such purposes (Dai and Li, 2022). This could potentially offer novel insights into how regulatory inheritance shifts under different stress conditions. These extensions underscore the versatility of our framework and highlight the need for continued development of statistical methodologies to harness the full potential of mother-daughter cell data.

## Supplementary Materials

ODEinherit is publicly available as an R package at https://github.com/WenbinWu2001/ODEinherit. The repository also includes the processed time series data used in this study. For network estimation, the codebase contains a computationally optimized version of KODE, originally implemented in MATLAB and generously provided by Lexin Li.

## Acknowledgments

We thank Lexin Li for providing the original code for KODE, which we further adapted into R for this work. We also thank Didong Li, Jingyi Jessica Li, Eardi Lila, Shirley Mathur, Ali Shojaie, Chang Su, Kevin Wang, and Yimin Zhao for the discussions that helped the ideas in this paper.

## Data Availability Statement

The processed time series data used in this study are publicly available at https://github.com/WenbinWu2001/ODEinherit under the data folder.

