## supplements for "Measuring Regulatory Network Inheritance in Dividing Yeast Cells Using Ordinary Differential Equations"

In this Appendix, we provide additional details regarding our methodology, simulations, and data. Section S1 describes the protein variables in our dataset and details our data preprocessing steps. Section S2 provides more details on the estimation procedure of KODE, hyperparameter tuning, and the detailed algorithm for network pruning. Section S3 elaborates on the random network generation and the ROC curve estimation.

### S1 Details of Yeast Cell Data

Section S1.1 provides details about the proteins in our dataset. Section S1.2 provides details about how the data was preprocessed after the features were extracted from the live-cell microscopy experiment.

#### S1.1 Protein Variables in the Analysis

The six measured proteins and their corresponding fluorescent markers are as follows. The first protein is Cdc10, labeled with mCyOFP1, which is tracked as the standard deviation of its fluorescence at the cell periphery; this serves as a marker for the septin ring and indicates cell cycle stages. The remaining proteins include Protein Stb3 (labeled with mTFP1), Protein CLB5pr-newNClb2-NG (labeled with mNeonGreen), Protein Whi5 (labeled with mKOk), Protein Xbp1 (labeled with mRuby3), and Protein Tup1 (labeled with mNeptune2.5), all of which are measured by their nuclear concentrations. These proteins monitor key cell cycle activities.

We use our previously developed segmentation method based on convolutional neural networks called FIEST (Ramakanth et al., 2024) to precisely segment cells and computationally track individual cells across the experiment. The resulting time-course data contains labels for time-of-birth and mother-daughter affiliations across all divisions. Currently, our dataset is limited to six proteins due to constraints in real-time live-cell microscopy (i.e., imaging), driven by biotechnological challenges such as spectral overlap and binding specificity. For instance, using too many fluorescent markers can compromise the reliable detection of individual color channels.

#### S1.2 Details of Data Preprocessing

During the preprocessing, cells with more than 50% missing values in the trajectories for any variable were excluded. Furthermore, the raw trajectories exhibit high-variance noise. To filter out these noise signals, we projected each trajectory into the principal component space using Functional Principal Component Analysis (FPCA), which was applied to the trajectories of cells from all generations for each variable. We retained the leading principal components to cumulatively explain at least 99% of the variance. Following FPCA, we performed local polynomial regression of degree 3 with a bandwidth of 1 on each individual trajectory to further denoise and interpolate it, achieving a fivefold resolution on the observed time grid. Specifically, an original time grid from 1 to 48 with a step size of 1 is now interpolated to a step size of 0.2. Then, each trajectory is centered to have a zero mean.

To deploy the above procedure, notably, the daughter trajectories are incomplete since their pre-birth portions are unobservable, which prevents a direct application of FPCA. To address this, we developed a correlation-based approach to impute the missing portions of the daughter trajectories. Based on our empirical observation that mother and daughter trajectories are positively correlated, for each variable, we impute the pre-birth portion of the daughter trajectory using its Pearson's correlation with the mother trajectory after the birth (here, the correlation is computed by treating the trajectory values separately rather than as functional data). It is important to note that this imputation is solely for enabling the application of FPCA and is not used in subsequent inheritance estimation. The algorithm for this imputation procedure is detailed below in Algorithm 1.

---

**Algorithm 1:** Imputation of Daughter Trajectories Before Birth

---

**Input** : Observed daughter trajectory $y^{(D)}_{[n_0:n]}$ and observed mother trajectory $y^{(M)}_{[1:n]}$ for any variable $j \in \{1, \ldots, p\}$, where $n_0 \geq 2$ is the daughter's birth time.

**Output:** Imputed pre-birth portion of the daughter trajectory, $\widehat{y}^{(D)}_{[1:(n_0-1)]}$.

```
// Step 1: Smooth raw trajectories
```

Apply local polynomial regression (degree 2, bandwidth 2) to $y^{(M)}_{[n_0:n]}$ and $y^{(D)}_{[n_0:n]}$, obtaining smoothed trajectories $\widetilde{y}^{(M)}_{[n_0:n]}$ and $\widetilde{y}^{(D)}_{[n_0:n]}$.

```
// Step 2: Fit a linear relationship
```

Regress $\widetilde{y}^{(D)}_{[n_0:n]}$ on $\widetilde{y}^{(M)}_{[n_0:n]}$ to estimate the intercept $\widehat{\alpha}$ and slope $\widehat{\beta}$.

```
// Step 3: Predict pre-birth daughter trajectory
```

Predict the pre-birth daughter trajectory as:

$$\widehat{y}^{(D)}_{[1:(n_0-1)]} = \widehat{\alpha} + \widehat{\beta} \cdot y^{(M)}_{[1:(n_0-1)]}.$$

// Step 4: Adjust for continuity at $t = n_0$

Update the prediction to ensure continuity at the daughter's birth:

$$\widehat{y}^{(D)}_{[1:(n_0-1)]} \leftarrow \widehat{y}^{(D)}_{[1:(n_0-1)]} + \left(y^{(D)}_{[n_0]} - \widehat{y}^{(D)}_{[n_0]}\right),$$

where $\widehat{y}^{(D)}_{[n_0]} = \widehat{\alpha} + \widehat{\beta} \cdot y^{(M)}_{[n_0]}$.

**return** $\widehat{y}^{(D)}_{[1:(n_0-1)]}$.

---

### S2 Details of Methods

Section S2.1 provides additional details on how KODE works, as ODEinherit uses KODE for network estimation prior to deploying our pruning procedure. Section S2.2 discusses the kernels we use in this paper. Section S2.3 provides additional details on how our pruning procedure works.

#### S2.1 Details of KODE Estimation

In this section, we provide further details of the KODE estimation procedure (Dai and Li, 2022), which is based on a two-step collocation estimation framework. In the first step, smoothing estimates of the observed trajectories are obtained using (3). In the second step, the functionals $F_j$'s are estimated by solving the penalized optimization problem (4), which is reformulated for computational efficiency as follows: for $j \in \{1, \dots, p\}$, solve

$$\min_{\theta_{j0},\theta_j,F_j} \frac{1}{n}\sum_{i=1}^{n}\left\{y_{ij} - \theta_{j0} - \int_0^{t_i} F_j(\widehat{z}(t))dt\right\}^2 + \eta_{nj}\Big(\sum_{k=1}^{p}\theta_{jk}^{-1}\|\theta_{jk}F_{jk}\|_{\mathcal{H}}^2\Big) + \kappa_{nj}\Big(\sum_{k=1}^{p}\theta_{jk}\Big)$$
$$\text{such that:} \quad \theta_{jk} \geq 0, \quad k \in \{1, \dots, p\}, \tag{11}$$

where $\eta_{nj}, \kappa_{nj} \geq 0$ are the tuning parameters. This problem is solved via an iterative alternating optimization approach, updating each component of $(\theta_{j0}, \theta_j, F_j)$ while holding the others fixed in each iteration. Notably, when $\theta_{j0}$ and $\theta_j$ are fixed, the solution $\widehat{F}_j$ takes the form of a finite linear combination of kernel products, owing to a generalization of the representer theorem. This allows $F_j$ to be estimated through a finite set of parameters.

The procedure involves four tuning components. In step 1 mentioned in the main text (see (3)), $\lambda_{nj}$ regularizes the smoothing estimates and is chosen by generalized cross-validation (GCV). In step 2, when solving the reformulation in (11), the kernel choice and bandwidth selection are detailed in the next section. $\eta_{nj}$ imposes a Ridge-type penalty to $F_j$ in the RKHS $\mathcal{H} = \{1\} \oplus \sum_{k=1}^{p}\mathcal{H}_k$ and is tuned by GCV. $\kappa_{nj}$ controls sparsity in $\theta_j$ through a Lasso regularization and is selected by ten-fold cross-validation.

#### S2.2 Kernels Used in KODE

In our implementation of KODE, we consider the following two types of kernels.

- *Gaussian kernel*: The Gaussian kernel is defined as $K(x, x') = \exp\{-\frac{(x-x')^2}{2\sigma^2}\}$ for $x, x' \in \mathbb{R}$. We follow the strategy given by Mukherjee and Zhou (2006) and Yang et al. (2016) to choose the bandwidth $\sigma$. In particular, $\sigma$ is set as the median over all pairwise distances among all sample points. The RKHS generated by the Gaussian kernel contains infinitely differentiable functions (Lin and Brown, 2004), giving us a class of smooth functions.

- *First-order Matérn kernel*: The first-order Matérn kernel is defined as $K(x, x') = (1 + \frac{\sqrt{3}|x-x'|}{\nu}) \exp\{-\frac{\sqrt{3}|x-x'|}{\nu}\}$ for $x, x' \in \mathbb{R}$. The bandwidth $\nu$ is chosen to be 1. The RKHS generated by the first-order Matérn kernel contains once differentiable functions (Gneiting et al., 2010), giving us a class of more flexible functions.

#### S2.3 Pruning a Network From KODE

We detail the pruning procedure for the estimated network from KODE in Algorithm 2. While sparser networks can also be obtained by increasing the Lasso regularization in KODE, the selection consistency depends on sufficiently strong regulatory effects in true edges and negligible effects in non-edges (Dai and Li, 2022; Chen et al., 2017). These assumptions can be challenging to satisfy under model misspecification or in the RKHS space, where the functional estimands take more complex forms. Simply thresholding the number of selected edges in Lasso is suboptimal, as it requires prior knowledge of the network and provides little insight into the significance of selected regulators. Our approach provides an explicit quantification of the regulatory contribution from each of the six proteins while remaining computationally feasible. Empirically, this strategy demonstrates superior performance. However, in high-dimensional settings beyond the scope of this paper, imposing larger Lasso regularization likely remains a practical and effective choice.

**Algorithm 2:** Pruning the Estimated Network From KODE

**Input :** The estimated network $\{M_j : j \in \{1, \ldots, p\}\}$ and a pruning threshold $\tau \in [0, 1]$ (by default, 0.05).

**Output:** A pruned network $\{\widetilde{M}_j \subseteq M_j : j \in \{1, \ldots, p\}\}$.

Initialize the current network $\widetilde{M}_j \leftarrow M_j$ for $j \in \{1, \ldots, p\}$;<br>
**for** *variable* $j \in \{1, \ldots, p\}$ **do**<br>
    `// Prune the regulator set of each variable separately`<br>
    **repeat**<br>
        `// Evaluate goodness-of-fit of the current regulator set`<br>
        Compute $\widetilde{R}_j^2$ using $\widetilde{M}_j$;<br>
        **for** *each selected regulator* $k \in \widetilde{M}_j$ **do**<br>
            Remove this regulator from $\widetilde{M}_j$, and obtain $\widetilde{M}_{j,-k} \leftarrow \widetilde{M}_j \backslash \{k\}$;<br>
            `// Evaluate goodness-of-fit of the reduced regulator set`<br>
            Compute $\widetilde{R}_{j,-k}^2$ using $\widetilde{M}_{j,-k}$;<br>
            Compute the reduction as $\bar{\delta}_{j,-k} \leftarrow \widetilde{R}_j^2 - \widetilde{R}_{j,-k}^2$;<br>
        **end**<br>
        Find the "least important" regulator as $k^* \leftarrow \arg\min_{k \in \widetilde{M}_j} \bar{\delta}_{j,-k}$;<br>
        **if** $\bar{\delta}_{j,-k^*} < \tau$ **then**<br>
            Prune the edge from $k^*$ to $j$, and update $\widetilde{M}_j \leftarrow \widetilde{M}_j \backslash \{k^*\}$;<br>
        **else**<br>
            `//` $\widetilde{M}_j$ `is pruned to the simplest, prune the next variable`<br>
            **break**<br>
        **end**<br>
    **until**;<br>
**end**<br>
**return** $\{\widetilde{M}_j : j \in \{1, \ldots, p\}\}$

### S3 Details of Simulations

Section S3.1 provides additional details on how the simulated datasets were generated, how the initial conditions were set, and how we numerically computed the ODE estimates in the simulations. Section S3.2 provides additional details about how we generated random networks when creating a baseline in our experiment to assess the inheritance score. Section S3.3 provides additional details on how the ROC curves were plotted in Figure 3. Section S3.4 provides additional results about the network estimation based on our simulated data.

#### S3.1 Additional Simulation Setup

We provide more details not discussed in Section 4. In the implementation of KODE, without loss of generality, we treat the data observations as if they were observed on the standardized interval $\mathcal{T} = [0, 1]$. For example, if the data are observed on an evenly spaced interval with sample size $n$, then the observed time points are standardized to $\{1/n, 2/n, \ldots, 1\}$. We employ KODE to estimate the regulatory network of this system from the data observations. For simplicity, each trajectory is centered to have a zero mean. In step 1 of KODE, the smoothing estimates $\widehat{z}(t)$ are obtained using cubic smoothing splines with knots at each observed time point, where the smoothing parameter is chosen by GCV. In step 2, we use a Gaussian kernel for each variable for the additive system and a first-order Matérn kernel with $\nu = 1$ for each variable for the non-additive system. Numerical solutions of this system are obtained by the Euler method with step size $10^{-5}$ on $[0, 1]$.

We discuss how the initial conditions were set in our simulations.

- *Network estimation* (Section 4.1): In the additive ODE simulations, the initial conditions of each triplet are sampled independently from $\text{Unif}(-A^{(k)}, A^{(k)})$. In the non-additive ODE simulations, the initial conditions are sampled from $\text{Unif}(\{-20, -19, \ldots, 20\})$ for all variables, independent of all other variables.

- *Inheritance score* (Section 4.2): In the additive ODE simulations, the initial conditions are still randomly sampled for each simulation run. In the non-additive ODE simulations, we fix the initial conditions in these simulations to more clearly illustrate the performance of our inheritance score under model misspecification. Specifically, the initial conditions in the non-additive system are fixed at $(-3, 15, -20)$ and $(-1, 1, 0)$ for the two triplets throughout these simulations.

For the additive ODE simulations, when estimating the networks, the integrals in (4) and (5) are numerically computed on a fine grid on the interval $[0, 1]$ with step size 0.001.

For the non-additive ODE simulations, when estimating the networks, we obtain the numerical solutions using the ODE solver `deSolve::ode` in R with the `lsoda` integrator (Hindmarsh, 1983; Soetaert et al., 2010) with step size 0.01 on $[0, 100]$.

#### S3.2 Generating Random Networks of Equivalent Density

We describe how we generate the random networks introduced in Section 4.2, which were used to generate the results shown in Figure 5. For a given network $\{M_j : j \in \{1, \dots, p\}\}$, a random network of equivalent density is generated by sampling $|M_j|$ elements from $\{1, \dots, p\}$ for each variable $j$ so that the number of regulators of each variable remains unchanged.

#### S3.3 Estimating an ROC Curve

We provide additional details on how the ROC curves in Figure 3 were plotted. To estimate an ROC curve from a set of (FPR, TPR) pairs, we proceed in two steps following the strategy adopted by Lin et al. (2021). First, for each (FPR, TPR) pair, linear interpolation is used to estimate the TPR values over a fine grid of FPR values. Second, the median TPR value is computed at each FPR point on the grid to construct the ROC curve. The AUC is then calculated using the trapezoidal rule.

#### S3.4 Additional Network Estimation Results

We provide additional details on ODEinherit's network estimation performance, before and after pruning, for both the additive and non-additive ODE systems to supplement the results in Section 4.1. In the additive ODE system, there are 12 true edges among 36 possible edges. On average, our estimated network selects $(12, 16, 13)$ edges before pruning and $(12, 11, 10)$ edges after pruning, corresponding to noise levels of $(0\%, 3\%, 10\%)$, respectively. In the non-additive ODE system, there are 16 true edges among 36 possible

| System | Type | Noise level | | |
|---|---|---|---|---|
| | | 0% | 3% | 10% |
| Additive | Original $R^2$ | 1.00 | 0.85 | 0.81 |
| | Pruned $R^2$ | 1.00 | 0.87 | 0.83 |
| Non-additive | Original $R^2$ | 0.99 | 0.98 | 0.96 |
| | Pruned $R^2$ | 0.98 | 0.97 | 0.95 |

Table S1: $R^2$ values for the original and pruned estimated networks in simulations of additive and non-additive systems at various noise levels.

edges. On average, our estimated network selects $(27, 25, 27)$ edges before pruning and $(9, 7, 12)$ edges after pruning, corresponding to noise levels of $(0\%, 3\%, 10\%)$, respectively. Table S1 reports the $R^2$ values of the refitted trajectories using networks before and after pruning. We observe that pruning substantially reduces network density while preserving a similar level of explanatory power for the observed trajectories.
